# From stuff to things: Responses of neurons in macaque V4 to textures and objects

**DOI:** 10.1101/2024.02.20.581273

**Authors:** Justin D. Lieber, Timothy D. Oleskiw, Laura Palmieri, Eero P. Simoncelli, J. Anthony Movshon

## Abstract

Humans and monkeys can rapidly recognize objects in everyday scenes. It is not fully understood where this computation arises, but previous work suggests selectivity for object shape first emerges in cortical area V4. To explore the mechanisms of this selectivity, we generated images on a continuum between “scrambled” textures and photographic images, preserving the local statistics of the original image while discarding structural information about scene and shape. We defined this continuum by the size of the region in which statistics were measured (“pooling region”). On average, neurons responded equally to photographic images and their scrambled counterparts. However, neurons exhibited a greater dynamic range of response to photographs. As a result, neuronal populations in V4 could more reliably discriminate between photographs than between scrambled images. Responses to partially scrambled images were more similar to responses to fully scrambled images than to photographs, even for perceptually subtle changes, and only began to resemble responses to photographs for pooling regions roughly half the size of typical V4 receptive fields. These same patterns emerged in an image similarity metric designed to predict human judgements of image degradation. Finally, V4 object selectivity showed dynamics that grew slowly and persisted following response offset, suggesting this signal may arise from recurrent mechanisms.

**Significance Statement:** Object recognition is a primary goal of visual processing, but is too complex to be computed by early visual areas. To evaluate where object-selective signals emerge, we recorded the responses of individual neurons in mid-level area V4 as macaque monkeys viewed images of objects (“things”) and of matched textures (“stuff”). Object selective V4 responses emerged in two phases. Short-latency responses discriminate objects from one another, but the distinction between objects and textures arrived later. This late-emerging signal was sensitive even to small image degradations, as is human perception. Our results show how V4 initiates the brain’s representation of visual objects.

## Introduction

Humans and monkeys are adept at recognizing objects in everyday scenes. The neural substrate for object recognition is a series of computations in the ventral stream of visual cortex (Ungerleider and Mishkin, 1982; Mishkin et al., 1983; Goodale and Milner, 1992; Logothetis and Sheinberg, 1996; DiCarlo et al., 2012; Kaas et al., 2022). This stream consists of a series of hierarchically connected visual areas, beginning in primary visual cortex (V1), continuing through higher cortical areas V2 and V4, and culminating in inferotemporal cortex (IT). Population responses in IT can successfully discriminate visual objects (Pasupathy and Connor, 2002; Rust and DiCarlo, 2010) and match human performance on object recognition tasks (Majaj et al., 2015). However, the intermediate computations that build these object selective responses are only beginning to be understood.

One of these intermediate computations is a growing selectivity for the characteristic features of natural images. Neurons in areas V2 and V4 are responsive to statistical regularities present in naturalistic texture images, unlike neurons in area V1 (Freeman et al., 2013; Okazawa et al., 2016; Ziemba et al., 2016, 2018, 2019, 2024). In V2, this modulation emerges 60-80 ms after response onset (Freeman et al., 2013; Okazawa et al., 2016; Ziemba et al., 2018); in V4 the modulation is both stronger and more delayed (Okazawa et al., 2016; Lee et al., 2024, 2025). In V4, responses are more strongly modulated by photographic images than by synthetic textures with matched statistics (Rust and DiCarlo, 2010; Movshon and Simoncelli, 2014; Long et al., 2018; Kramer et al., 2023; Kim and Pasupathy, 2024).

Another proposed intermediate computation is spatial pooling. Ventral stream neurons integrate their preferred visual inputs over regions of a certain size. Neuronal responses may compute a statistical summary of image features in this region, discarding information about their precise spatial arrangement (Lettvin, 1976; Wilkinson et al., 1997; Parkes et al., 2001; Pelli et al., 2004; Pelli and Tillman, 2008; Balas et al., 2009; Greenwood et al., 2009; Freeman and Simoncelli, 2011; Rosenholtz et al., 2019; Ziemba and Simoncelli, 2021; Broderick et al., 2025). This *pooling region* determines a critical distance: positional information within that region is lost; beyond that region, it is preserved. Pooling seems to reflect the growth in neuronal receptive fields along the ventral stream (Gross et al., 1972; Gattass et al., 1981, 1988). Spatial pooling may explain why V4 responses show invariance to translations of visual objects (Pasupathy and Connor, 1999; Rust and DiCarlo, 2010; Nandy et al., 2013; Sharpee et al., 2013; El-Shamayleh and Pasupathy, 2016), and why these invariances are stronger in IT (Rust and DiCarlo, 2010; Hong et al., 2016), where receptive fields are larger.

To evaluate how object-selective signals emerge, we recorded the responses of individual neurons in V4 as macaque monkeys viewed images of objects (“things”) and of matched textures (“stuff”). We selected a set of photographs of everyday scenes, then “scrambled” their features so that global spatial information was preserved but local spatial information was lost. By adapting a texture synthesis algorithm (Portilla and Simoncelli, 2000), we synthesized images that matched statistics in localized regions. By varying the region sizes, we generated images that transition smoothly between photographs and textures.

Classification analyses using the responses of V4 populations showed that photographic image responses were substantially easier to discriminate than partially scrambled images, and that this increased discriminability began at a critical size that was smaller than the size of typical V4 receptive fields. This pattern of neural activity was qualitatively explained by DISTS, an quality assessment measure built to capture human sensitivity to image distortions. The neural dynamics of discriminability showed that V4’s sensitivity to photographic structure emerged late relative to the onset of neural activity, and persisted well after stimulus offset, suggesting it arises from recurrent or feedback processes.

## Materials and Methods

### Image generation

We chose a core set of 20 photographic images from two databases. Half were taken from the UPenn Natural Image Image Database: photographs taken at a baboon habitat in Botswana (Tkačik et al., 2011) that approximate the evolutionary context for primate vision. The other half were taken from the Reachspace Database, which features everyday objects in their appropriate context (Josephs et al., 2021). These images of objects on natural backgrounds are similar to those typically used to train and evaluate deep neural network models of area V4 (Yamins et al., 2014). All source images were cropped and resolution adjusted to contain objects centered within a 1280×1280 pixel frame. From each source image, we then cropped four overlapping 512×512 subimages, centered at locations +/-56 pixels horizontally and vertically displaced from the center of the source image. In the experiment, images were suitably vignetted and presented on a screen at a resolution of 80 pixels per degree. Therefore, when presented, these “shifts” of the underlying parent image correspond to an angular distance of 1.4 deg.

To generate scrambled images, we used an adaptation of a texture synthesis model that measures and synthesizes images with statistics that are matched within localized pooling subregions (Freeman and Simoncelli, 2011) (https://github.com/freeman-lab/metamers). For this study, we arranged these pooling regions as a square grid of smoothly overlapping fields that tiled the image. Within each region, we measured a set of texture statistics (Portilla and Simoncelli, 2000). Specifically, we processed the image by convolving it with a bank of 16 oriented filters (4 orientations, 4 scales, 2 phases represented as real and imaginary components), computed with periodic boundary conditions. To create “simple-cell-like” and “complex-cell-like” channels, we computed the linear and amplitude responses of each filter, respectively. We then computed the pairwise product of these channels for different pairs of positions, orientations, and scales. Finally, we computed weighted averages of these products within each pooling region, resulting in a set of covariance statistics.

We synthesized new “scrambled” images by generating an initial Gaussian white noise image, then iteratively adjusting the pixels to match the measured texture statistics within each subregion. To prevent boundary artifacts from appearing within the synthesized images, we defined a circular vignette with diameter 492 pixels (6.15 deg) and raised cosine edges spanning 25 pixels (0.31 deg). Between iterations of the synthesis we blended the edge regions of the image (using blending fractions governed by the vignetting function) with the pixel values of the original photographic image. After synthesis was complete, the images were cropped with this same circular vignette to remove these “photographic” regions, such that only “scrambled” portions of the image remained.

For each of the four “shifted” 512×512 subimages, we synthesized new images based on 1×1, 2×2, 3×3, 4×4, and 6×6 grids of statistical pooling windows. When presented to the subject, these pooling regions of these grids subtended 6.4, 3.2, 2.1, 1.6, and 1.1 deg, respectively. We also included the original (unscrambled) images, giving a total of 6 distinct conditions for each original photographic subimage. We refer to the set of 24 images derived from an original photograph (4 shifts x 6 pooling window sizes) as an “image family.”

### Experimental procedures

Experimental procedures for monkeys conformed to the National Institute of Health *Guide for the Care and Use of Laboratory Animals*, and were approved by the New York University Animal Welfare Committee.

We recorded eye position with a high-speed, high-precision eye tracking system (EyeLink 1000). The animal initiated each trial by fixating on a small dot (∼0.25 degrees wide), and maintained fixation within a window of 1-2 degrees. Fixational drift was small, typically staying within a 0.5 deg of the fixation point (mean std dev. of eye position per trial = 0.24 deg, see *Materials and Methods: Eye movement analysis*). Images appeared for 200 ms, with a 200 ms inter-stimulus interval, and were blocked in 3-8 consecutive presentations, after which the animal received a juice reward.

### Acute recordings

Under general anesthesia, we implanted one male *macaca mulatta* (Monkey M) with a titanium head post and recording chamber over area V4. After recovery, we recorded single unit activity within V4 using both single site electrodes (FHC) and a linear microelectrode array (Plexon S-probe, 64 channels with 50 µm spacing, over 3.2 mm total) while the animal stared at a small fixation dot. We identified the location of the lunate sulcus, then recorded in surface V4 anterior to the sulcus.

For single-site electrode recordings, we recorded all cells we found that were well-isolated and could be reliably driven by the image set. For linear array recordings, we inserted the probe into cortex deeply enough that visual stimuli drove multi-unit responses on most channels. The centers of neuronal receptive fields lay between 1 and 12 degrees from the center of gaze (median=6.1 deg). We isolated single units from multi-site recordings using KiloSort 2.5 (Steinmetz et al., 2021) followed by manual curation of well-isolated spikes (Phy, https://phy.readthedocs.io/). Cells were only analyzed when recordings lasted long enough to get at least 4 repetitions to all image stimuli (50/82 for single-site electrodes, 117/123 for linear arrays), and when the evoked firing rate to all image stimuli was at least 2 spikes/second (44/50 for single-site electrodes, 79/117 for linear arrays), for a total of 123 analyzed units from Monkey M.

### Chronic array recordings

Under general anesthesia, we implanted a 96 electrode “Utah” recording array (Blackrock) in area V4 of one male *macaca nemestrina* (Monkey B). We took area V4 to lie on the prelunate gyrus between the lunate and superior temporal sulci, dorsal to the tip of the inferior occipital sulcus. Electrodes shanks were 1 mm long and were arranged in a square grid (spacing 0.4 mm).

We recorded neural responses to our stimulus set over a period of five weeks, during which we collected 15 recording sessions. For a subset of these channels, we mapped receptive field centers; all lay between 1 and 2 degrees of eccentricity. We isolated single-units from these recordings using Mountainsort 5 (Chung et al., 2017) with SpikeInterface (Buccino et al., 2020), followed by manual curation of well-isolated spikes (as above). Because recording sessions on an electrode from consecutive days could potentially contain the same units (Dickey et al., 2009), we chose to use a subset of the recording sessions for full analysis. Specifically, for each electrode, we identified every session with a well-isolated single unit, then characterized the ability of those units to discriminate image responses from blank trial responses. We then selected the single day of recording that had the highest level of stimulus vs. blank discriminability. After removing electrode channels that never showed responsive single units, this resulted in a final population of 88 single-unit recordings from Monkey B.

### Subpopulations based on receptive field eccentricity

For a subset of analyses in this paper, we split our neuronal population into 4 non-overlapping subpopulations, each covering a distinct subset of the range of receptive field center positions. The first of these groups consisted of the 88 single-unit recordings from Monkey B. We defined the median receptive field center of this group at 1.5 degrees eccentricity.

To create the remaining groups, we first ordered recordings from Monkey M by their estimates of receptive field center eccentricity. We chose two partition values (4.5 and 7.5 degrees) that split these recordings into three subsets of approximately equal size (40, 46, and 36 neurons, from foveal to peripheral) and progressing eccentricity (3.2, 6.4, and 9.2 degrees, respectively). Population analyses within these four subgroups were done on subsamples of 18 neurons, as described below.

### Eye movement analysis

For each image presentation, we quantified the stability of eye position by measuring its standard deviation, defined as the square root of the summed variance of the horizontal and vertical eye positions over the course of the image presentation and following inter-stimulus interval (400 ms). We computed mean standard deviation of eye position by first averaging over all stimulus presentations within a recording session, then across all recording sessions. Mean standard deviations were similar for both animals (Monkey M=0.24, Monkey B=0.25).

Within each recording session, for each individual image, we ordered its repeated presentations by their standard deviation of eye position. We then split these in half into “low” and “high” movement groups. For sessions with odd numbers of trials, the middle trial was randomly assigned to either group. We then repeated our modulation and classification analyses (see below) with each group’s subset of trials.

### Firing rate analyses

For all cells, we computed the average firing rate on each trial in an interval between 50 ms and 400 ms after stimulus onset. When comparing or combining data across neural populations, we computed a normalized firing rate by dividing all individual trial firing rates by the average firing rate response to all non-blank stimulus conditions.

### Stimulus-dependent fractional variance

The visually evoked firing rate responses of V4 neurons varied across both repeated presentations of the same stimulus, as well as different presentations of different stimuli. To quantify the extent to which differences in stimulus responses were reliable, we computed the fraction of overall variance that could be explained by differences in stimuli. Specifically, we computed total variance as the variance in firing rate responses across all trials, and we computed stimulus-dependent variance as the variance across each stimulus’s trial-averaged firing rate. Stimulus-dependent fractional variance was taken as the ratio of stimulus-dependent variance to total variance.

### Modulation index

We computed a modulation index for individual cells as a standardized measure of the signed strength of photographic vs. scrambled firing rates. We computed this index in three different contexts: 1) for each neuron’s response to each image family, 2) for each neuron overall, and 3) for each image family. In each case, the index (as shown in Figure 2A) is the difference divided by the sum of the average firing rates (over all image shifts) for photographs and scrambled images, respectively. To compute a modulation index for each image family, we first averaged firing rates across all neurons, then computed the modulation index as normal.

### Image family ranking

We sought to visualize whether the image families that most strongly drove neurons were also the families most strongly modulated by photographic structure. Our first thought was to simply rank each neuron’s preferred image families, but we were concerned that trial-to-trial fluctuations in firing rate might lead to overfitting that obscured the true magnitude of photographic image modulation. We computed a cross-validated ranking of image families by splitting the image shift conditions into training sets (2 diagonally opposite shift conditions) and testing sets (2 remaining shift conditions). We averaged firing rates over training set shift conditions and over photographic and fully scrambled conditions, then used these rates to rank image families. We then used the test set to compute average firing rates for the photographic and scrambled conditions, and sorted those values using the training set rankings. The rank order plots in Figure 3 are the average of two rank order plots made from the two possible partitions of training/testing data.

### Rate-difference correlation

For each neuron, we quantified whether the image families that most strongly drove the cell were also the image families that most strongly preferred photographic structure. To compute an image family’s evoked firing rate, we averaged firing rates over all shifts and over both photographic and fully scrambled conditions. To compute each image family’s difference between photographic and scrambled image responses, we first averaged firing rates over all shifts, then computed the difference. We then computed each neuron’s Pearson correlation between family-evoked rates and family-evoked differences.

### Photographic-scrambled correlation

For each neuron, we computed the correlation across families between its response to photographic images and its response to scrambled images. To compute a family’s evoked firing rate, we averaged firing rates over all trials and all shifts. We then computed the Pearson correlation between average photographic rates and average scrambled rates.

### Single cell discriminability (d’)

We quantified the ability of single neurons to discriminate between two stimulus classes as d’, a standard measure of discriminability:

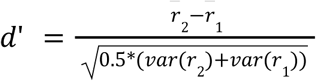

where *r*_1_ and *r*_2_ are the collection of single-trial firing rates for each class. To measure the ability of neurons to discriminate between photographic and scrambled images (Figure 3), we combined all the photographic and scrambled image trials from all image families into the two classes, then computed d’ values. For family vs. family discrimination (Figure 6A), we computed each neuron’s d’ for each pairwise comparison of two image families (either for photographic images, or for scrambled images), then averaged all unique family pairs into a single summary value.

### Classification with LDA

For a range of different tasks, we sought to quantify the ability of neural populations to discriminate between pairs of classes of stimuli. To this end, we implemented a cross-validated linear discriminant analysis (LDA). We first split our image set in half such that, for each image family, two diagonally opposite shift locations were used for model training, and the remaining two were used for testing.

For population classification tasks, we split our population into two matched sets of 105 neurons. We repeated this resampling process over 100 distinct splits of the full population into two distinct subpopulations. For each of these subpopulations, for each image, we constructed pseudo-trials by randomly selecting a single trial from each neuron. We repeated this process to create 500 pseudo-trials (with replacement) for each image.

We then used LDA to learn the weighted sum across neurons that best separated the two classes:

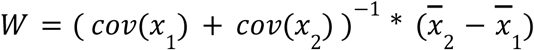

where *x*_1_ and *x*_2_ are matrices of training set firing rates corresponding to the two classes, respectively, with columns of neurons and rows of pseudo-trials of all images in the class, and *W* is the vector of resulting weights on the neurons. We then applied these weights to the testing data to compute a single scalar value for each trial, and computed discriminability (d’) using these scalar values.

To compute final discriminabilities, we averaged over both permutations of the trial splits into training and testing image sets, over both disjoint neural populations within each population resampling, and over all population resamplings. To compute error bars on performance values, we measured sample variances across the disjoint pair of neural populations, averaged sample variances over the resampled populations, then took the square root to find the standard deviation.

We trained classifiers for four distinct discrimination tasks:

*1) Stimulus vs. blank*: We trained classifiers to separate 1) a combined set of all image conditions (all image families, all scrambling conditions, and two shifts per training/testing set) and 2) blank trials. Blank trials were split in half into training and testing sets.
*2) Family-specific scramble vs. structured*: For each image family, we trained a classifier to separate fully scrambled images from either partially scrambled conditions or from the original photographic images. We averaged performance across all image families to get final discriminability values (Figure 4A, white points).
*3) Family-general scramble vs. structured*: For family-general discrimination (Figure 4A, grey points), we trained a single discriminant, shared across families, between 1) the full set of scrambled images and 2) similar combined sets of either partially or photographic image conditions.
*4) Pairwise family*: For each scrambling condition, we identified every possible pair of two image families. We trained separate discriminants for every pairwise condition. To get a final discriminability for each scrambling condition, we averaged performance over all pairs of families (Figure 6B). Additionally, to quantify whether these classifiers could generalize between photographic and scrambled images, we took the discrimination axes learned from fully scrambled image training set responses, and applied them to the corresponding photographic testing set responses. This is the “generalization” classification performance in Figure 6B.

### Dynamics of discriminability

In the standard analysis for all four tasks, sensitivities were computed using average firing rates computed between 50 and 400 ms following stimulus onset. For the dynamics analysis, discriminabilities were computed in a series of 50 ms bins, spaced every 5 ms.

We defined the latency of each task as the time it took to reach half the maximum value of its peak performance. To compute error bars on latency, we measured the latency of the upper and lower error bars, using the same half-maximum criterion. We defined the difference of these error latencies, divided by two, as the error value.

### Perceptual and neural distance metrics

We asked whether the patterns of response we observed in V4 could be explained by computational models of perceptual image similarity that are used to quantify the perceived difference between an original image and a corrupted counterpart (e.g. to measure the quality of a lossy file format). We computed pairwise distances for both the V4 population response and for a set of image similarity metrics. In all cases, we only computed distances within sets of images of the same family and the same shift (an original photographic image and its partially or fully scrambled counterparts). Each of these sets had 6 images, which resulted in 30 total comparisons per “shifted” photographic image. Over 20 image families and 4 shifts per family, this resulted in 2,400 total distances.

### Neural distance

We first split our neural population into two matched subpopulations of 105 neurons. We repeated this resampling process over 100 distinct splits. Within each subpopulation, for each image we found a 105-neuron vector of averaged, normalized firing rates. We then computed the Euclidean distance of each pair of vectors, and normalized that distance by the number of neurons.

### Image similarity metrics

We used 3 standard image similarity metrics to compare different images: RMS pixel distance, structural similarity (SSIM (Wang et al., 2004)), and Deep Image Structure and Texture Similarity (DISTS (Ding et al., 2022)). For RMS pixel distance, we first normalized each image so that 0 and 1 corresponded to minimum and maximum luminance. We then computed the RMS difference over pixels for each pair of images. We computed SSIM using the *ssim()* function provided in Matlab, with all exponents set to 1. For the DISTS metric, we used a Matlab implementation of the algorithm provided by the authors (https://github.com/dingkeyan93/DISTS).

As a summary statistic, we computed the Pearson correlation between each image similarity metric and the corresponding neural distance. Correlations were averaged over all subpopulations. To compute error bars, we measured the sample variance across each disjoint pair of neural populations, averaged sample variances over the resampled splits of the population, then took the square root to find the standard deviation.

## Results

### Creating a continuum from scrambled textures to photographic images

We created a set of images that smoothly varied from fully scrambled textures to original photographs. To create this set, we first selected 20 photographic images from two natural image databases. Half were taken from the “Botswana” image set, a collection of evolutionarily relevant photographs taken at a primate nature reserve (Tkačik et al., 2011). The other half were taken from the “Reachspace” image set (Josephs et al., 2021), containing scenes of man-made objects in their typical context. From each of these images we extracted 4 discrete image “shifts” (Figure 1A), which contain similar (but not identical) content. For each of these shifted images, we synthesized a new set of images with matching texture statistics. To create varied “levels” of scrambling, we parametrically varied the size of the local pooling regions in which we measured texture statistics (Figure 1B). Images synthesized with smaller pooling regions were more similar to their progenitor photographs, and those synthesized with larger pooling regions were more distinct. Six scrambling levels were chosen to cover the range, from the original photographs to their globally scrambled counterparts, in approximately equal perceptual increments (Figure 1C). Counting all six shifts and four pooling region sizes, each original photograph created an image “family” containing 24 images.

**Figure 1.**
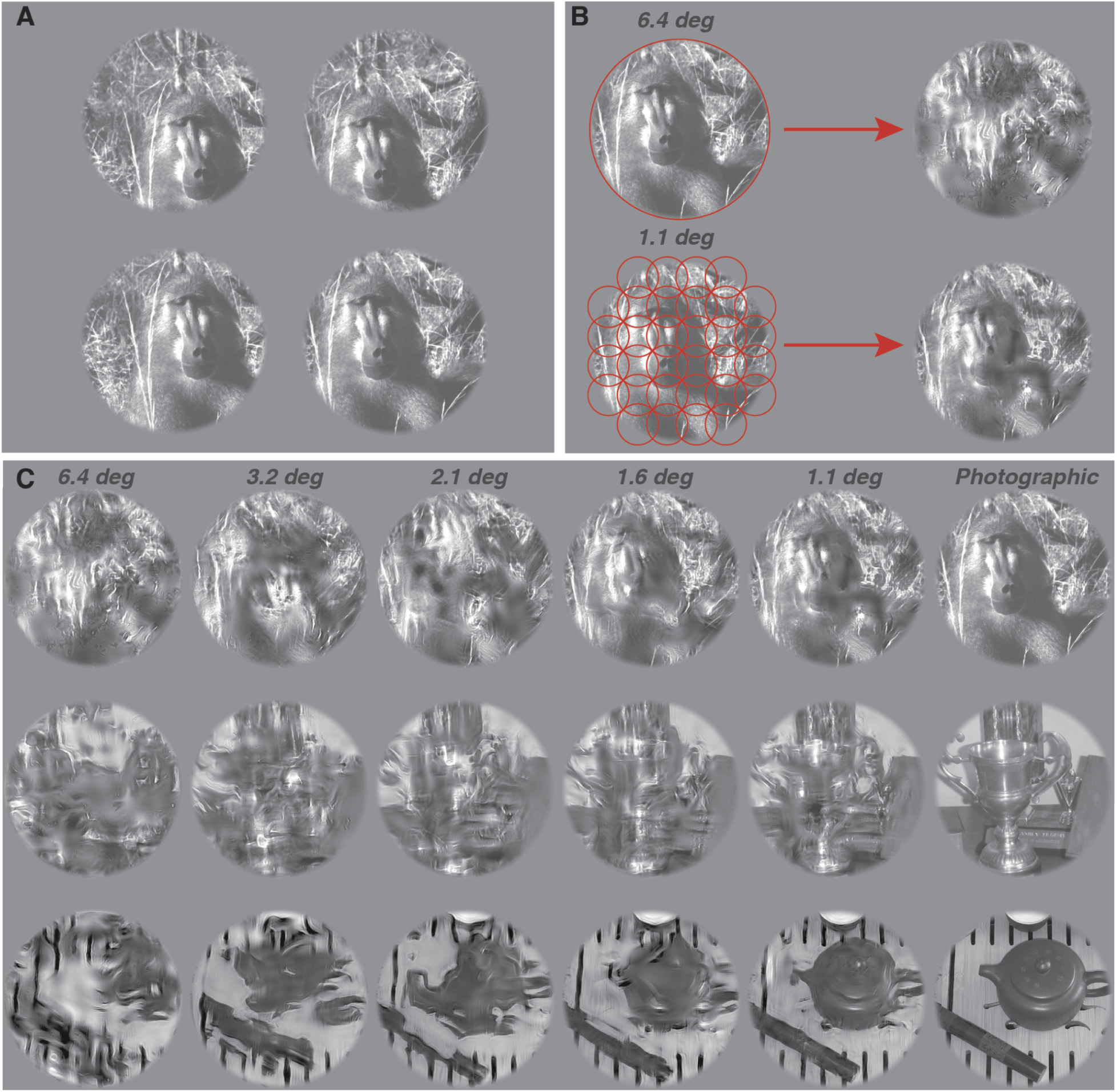
Creating a continuum from scrambled textures to photographic images. A) We cropped four distinct shifted subimages from each source image. These subimages were overlapping, and contained the same central features. B) We used the Portilla-Simoncelli texture model to scramble image features within local pooling regions (red circles). When a single pooling region covered the image (top) the content of the image was fully scrambled. When multiple smaller pooling regions covered the image (bottom), features within each pooling region were scrambled locally, and the image retained some of the global structure of the original. C) By varying the size of the pooling regions used in the image synthesis algorithm, we created a continuum of images between scrambled textures and the original images. We used 5 pooling region sizes across our 6.4 degree diameter image: 6.4, 3.2, 2.1, 1.6, and 1.1 degrees. Images smoothly transitioned from scrambled textures (left) to original photographic images of objects (right).

Our goal was to measure how V4 neuronal responses are affected by spatial scrambling. To quantify these effects, we focused on two distinct comparisons: the detection of photographic structure (whether V4 responses could distinguish between photographic images and their fully scrambled counterparts) and the discrimination of base images (whether V4 responses could distinguish between two distinct photographic images, and how that capability was degraded by scrambling). These two approaches can be visualized using the array shown in Figure 1C: comparing photographic vs. scrambled images corresponds to comparisons of different columns, and comparing photographic images to each other corresponds to comparisons of different rows.

### V4 neurons respond heterogeneously to photographic and scrambled images

We recorded 211 single units in area V4 that responded to the presentation of photographic and scrambled images over their receptive fields (see *Materials and Methods* for selection criteria). Nearly all neurons were significantly more active in response to images than to blank trials (197/211 cells significantly modulated, p<0.05, permutation test). To quantify whether V4 neuronal responses were responsive to differences between images, we computed the fraction of response variance that was captured by stimulus-driven modulation, rather than by trial-to-trial noise (see *Materials and Methods*). Fractional variances were typically greater than permuted controls (median fractional variance 0.369±0.180 median absolute deviation (MAD), 193/211 neurons with p<0.05, permutation test) indicating that V4 responses were significantly modulated by our image set.

Next, we compared V4 responses to photographic and scrambled images taken from the same image family. These pairs of images are matched in their spatially averaged statistics, and differ only in that the constituent features of scrambled images have been moved to “unnatural” locations. Differences in responses to these two classes of images reflects the ability of V4 responses to detect the characteristic “natural” features of photographic images when many simpler, potentially confounding features are matched.

We defined a scrambling modulation index (MI) as the difference between firing rate responses to photographic and scrambled images, divided by their sum. Modulation varied across neurons and image families (Figure 2A) from strongly positive (photograph-preferring) to strongly negative (scramble-preferring). A typical V4 neuron did not prefer photographic images to scrambled images (median MI for individual neurons = −0.002±0.050 MAD). Across V4 neurons (Figure 2B) a similar number exhibited either significantly positive (56/211, p<0.05, permutation test) or significantly negative (59/211) modulation. Population responses to image families were never strongly positively or negatively modulated (smallest family MI = −0.06, largest MI=0.05).

**Figure 2:**
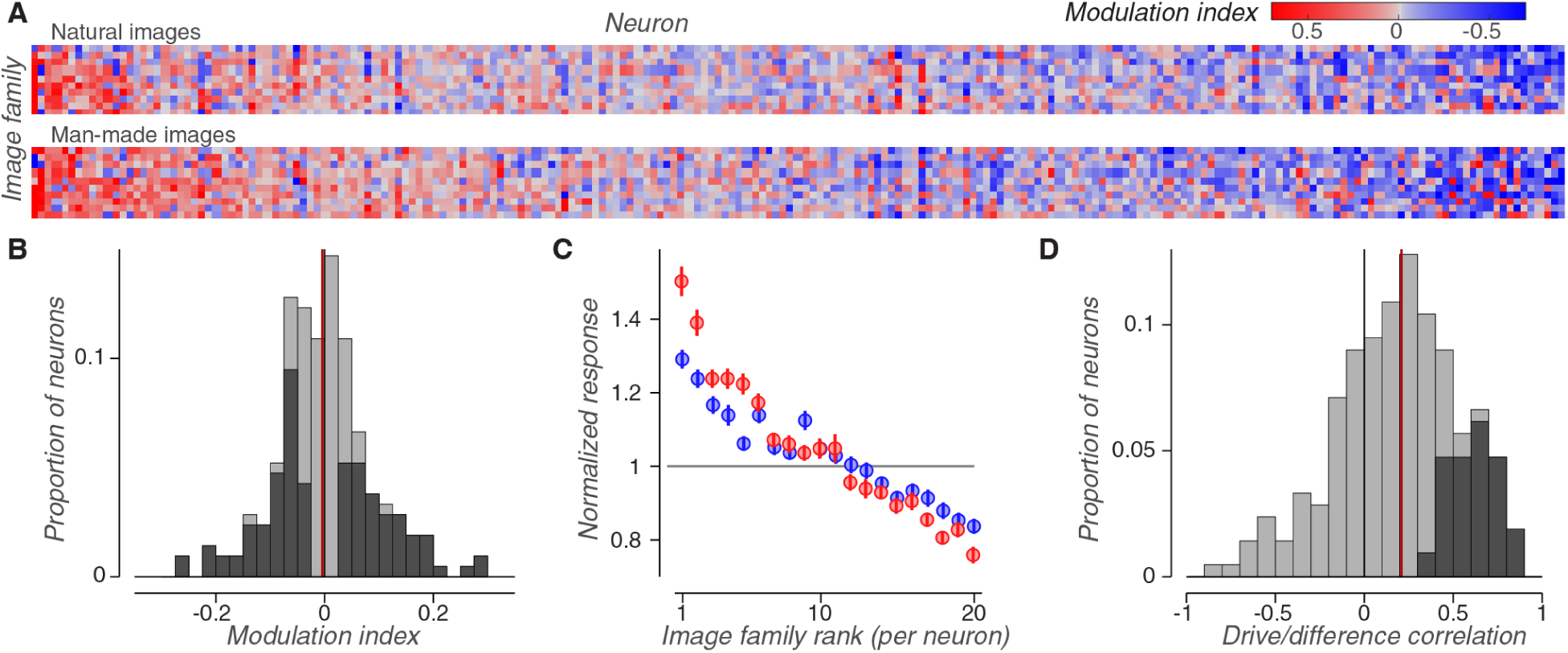
V4 neurons respond with greater dynamic range to photographs than to scrambled textures. A) Modulation indices of 211 neurons computed separately for 20 image families originating from two image sets: 10 from the “Botswana” images (top) and 10 from the “Reachspace” images (bottom). Within each set, both image families and neurons are ordered from highest to lowest modulation index, averaged over all points (top to bottom, and left to right, respectively). B) Distribution of the modulation indices of individual neurons, calculated on responses averaged across all image families. Neurons with modulation indices that are significantly different from 0 (p<0.05) are labeled in dark grey. Nearly as many neurons were significantly positively modulated (N=56, p<0.025) as significantly negatively modulated (N=59, p<0.025). The red line denotes the population average (MI=-0.004). C) Normalized firing rates plotted against the rank of image family drive. Ranks were computed separately for each neuron as a cross-validated measurement. Error bars denote standard errors across neurons and shifts. Strongly driven (highly ranked) families are modulated positively (preferring photographs) and weakly driven families are modulated negatively (preferring scrambled images). D) Distribution of individual neuron Pearson correlations between the average drive to an image family (averaged over both photographic and scrambled images) and that family’s scrambling difference (the difference between photographic and scrambled responses). Neurons with correlations significantly greater than 0 are labeled in dark grey (49/211, p<0.05). The red line denotes the median neuronal correlation (r=0.209). Most neurons had positive correlations, meaning that the image families that most strongly drove a neuron also tended to be positively modulated.

While the population tendency of V4 responses showed neither positive nor negative modulation, individual neurons showed idiosyncratic response patterns (Figure 2A). We reasoned that this heterogeneity in neuronal modulation might be explained by heterogeneity in neuronal tuning. For each neuron, we defined the “drive” for each image family as the average response to all of its photographic and fully scrambled images. Then, for each neuron, we ranked these families from strongest to weakest drive using a cross-validated procedure (see *Materials and Methods, Image family ranking*). On average, we found that high ranked families (those that evoked the strongest responses) were typically positively modulated, and that low ranked families (those driving the weakest responses) were negatively modulated (Figure 2C). For each neuron, we measured the correlation between each image family’s drive and its firing rate difference between photographic and scrambled images. These correlations were positive for most V4 neurons (Figure 2D, median Pearson correlation = 0.21±0.23 MAD, p<0.001, permutation test), validating the population trend visible in Figure 2C.

Photographic images typically drove cells to both their highest firing rates (for strongly driving families) and their weakest firing rates (for weakly driving families). We measured each neuron’s “dynamic range” for photographic and scrambled images as the variance of their respective normalized responses. Photographic image response variance (median σ^2^=0.098) was significantly larger than scrambled image response variance (median σ^2^=0.082) for most V4 neurons (152/211, p<0.001). This wider dynamic range for photographic images may explain previous reports of the superiority of V4 populations for discriminating photographic images over textures (Rust and DiCarlo, 2010).

### V4 responses discriminate between photographic and scrambled images

Both positively modulated and negatively modulated neurons can signal whether an image has photographic structure. We measured this informativeness using a standard measure of discriminability (d’) that we defined to be positive for both photograph-preferring neurons and for scramble-preferring neurons (see *Materials and Methods: Single cell discriminability (d’)*). These two subpopulations of V4 neurons were similarly effective at discriminating between photographic and scrambled images (Figure 3).

**Figure 3:**
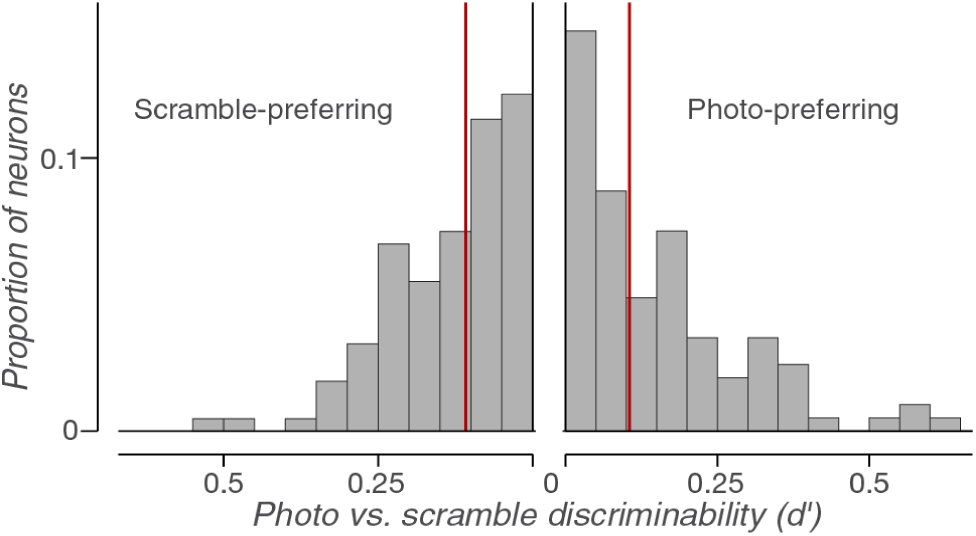
V4 neurons discriminate photographic scenes from their scrambled counterparts. Distributions of neuronal discriminabilities between paired photographic and scrambled images, either for neurons that prefer scrambled images (left) or photographic images (right). Discriminabilities are defined as positive for both populations. The red lines denote population medians (scramble d’=0.108, photographic d’=0.105).

We next measured how well neuronal population responses could work together to distinguish photographic image responses from their scrambled counterparts. We used linear discriminant analysis (LDA) to find the weighted sum of neuronal responses that best separated photographs and scrambled images. To ensure that our measurements of discriminability were not artificially inflated by noise or idiosyncratic responses to particular images, our LDA analysis was cross-validated across shifted images within each family (as in Figure 1A, see *Materials and Methods: Classification with LDA)*. Specifically, we used responses to two of the four shifts (training set images) of each photographic and scrambled image to find neuronal weights. We then computed d’ values by weighting responses to the two remaining image shifts (testing set images). The resulting d’ value is thus “translation-invariant”: it is only positive if the learned discriminant can successfully separate image translations that were not used to train the model. This process ensures that these classifiers do not rely on pixel-specific features of the images. Rather, they must rely on a more general response to “natural” image features.

We measured discriminabilities for each of the 20 image families individually, then took their average (“within-family discrimination”). Responses from small populations of V4 neurons (N=105) could reliably discriminate photographic and fully scrambled images (Figure 4A, white point on the right). We then extended this analysis to compare partially scrambled and fully scrambled image responses. If V4 neurons spatially pool their inputs, we would expect responses to partially scrambled images to resemble responses to photographs, provided that scrambling was within the region of spatial pooling. Contrary to this, we found that discrimination performance was strongly affected by partial scrambling (Figure 4A, white points), even for images where scrambling was confined to very small distances (64.5% lower for 1.1 deg pooling than for photographic images). We found this large difference in discriminability surprising, given that local scrambling only subtly changed how the images were perceived (cf. 1.1 deg vs. photographic in Figure 1C). This difference in discriminability began to rise at a pooling size of roughly 1.5 degrees. We take this as an estimate of the region over which image statistics are pooled by V4 neurons.

**Figure 4:**
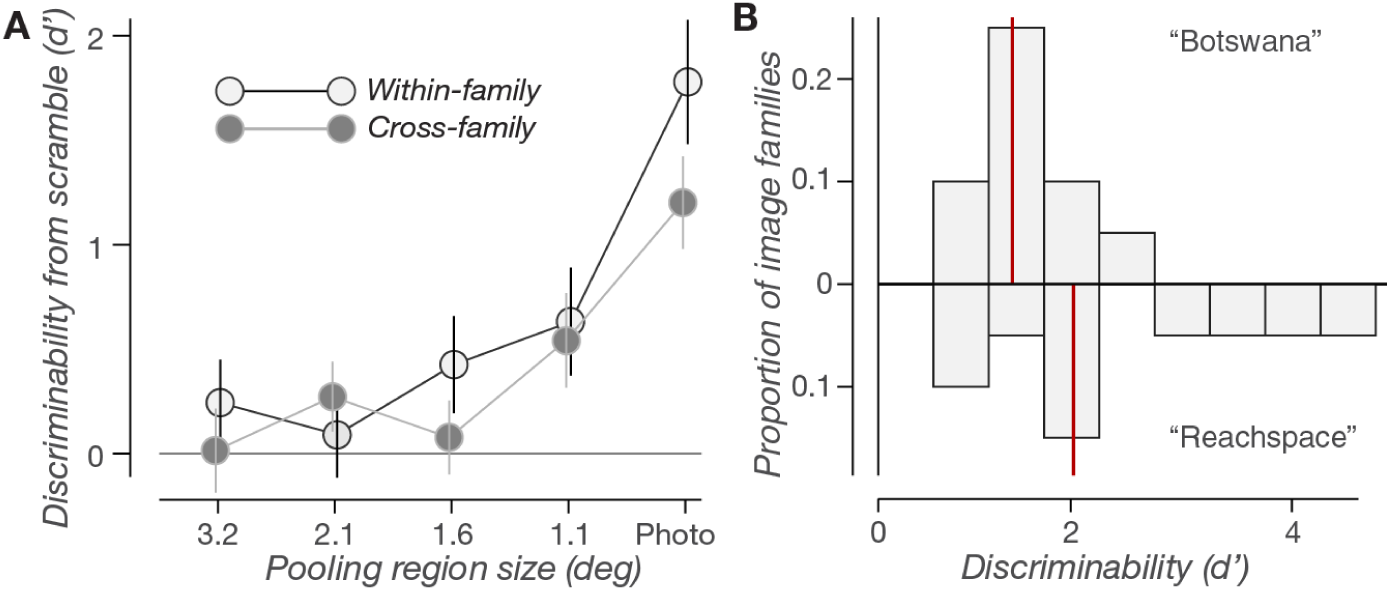
V4 populations discriminate photographic scenes from their scrambled counterparts. A) Discriminability of neuronal population responses (N=105 neurons) between fully scrambled image responses and either the original photographic images, or intermediate levels of scrambling. Discriminabilities were computed either for each image family then averaged (“within-family,” white points), or for all image families simultaneously (“cross-family,” grey points). Error bars show standard deviations across resampled neuronal populations. B) Two distributions of within-family discriminabilities, measured for either the 10 image families from natural scenes (“Botswana,” top) or the 10 image families from man-made scenes (“Reachspace,” bottom). Each discriminability is the median for the image family over resampled neuronal populations. The red lines denote medians over each image set (natural d’=1.21, man-made d’=1.77).

Considering individual image families (Figure 4B), we found that discriminabilities varied widely over a more than 7-fold range (min family d’=0.63, max=4.42). These differences were related, in part, to each image family’s parent database. Responses to photographs taken in “natural” environments were typically more difficult to discriminate (median d’=1.21, Botswana database) than those taken in man-made environments (median d’=1.77, Reachspace database).

Next, we investigated whether V4’s selectivity for photographic structure was invariant across different image families. Specifically, we asked whether we could identify a single readout axis that allowed the V4 population response to distinguish the full set of photographic images from all of their scrambled counterparts (“cross-family discrimination”). To this end, we retrained our photographic vs. scrambled classifiers using the full, combined set of image families (Figure 4A, grey points). These cross-family classifiers could reliably distinguish photographic and fully scrambled images, with only a 32% reduction in sensitivity compared to within-family classifiers. Like within-family classifiers, cross-family classifier performance was highly disrupted by partial scrambling (54% lower for 1.1 deg scrambled than photographic), and began to rise at roughly 1.5 degree pooling region sizes. Thus, V4 populations contain a single readout axis for photographic structure that is robust across different scenes.

Many of our recordings were taken from units with foveal receptive fields, whose responses were more likely to be affected by eye movements during trial fixations. While analysis of eye movement traces showed that fixational drift was small (mean std dev. of eye position per trial = 0.24 deg), we were concerned that even these small movements might still modulate neuronal responses. To check this, we split every stimulus’s repeated trials into “low” and “high” eye movement partitions (mean std. dev. of eye position for low partition=0.084, for high partition=0.408). Our modulation and classification analyses showed identical results using either low movement trials or high movement trials. We extended this split procedure to all other analyses in this study, and found similar results. We conclude that eye movements had no major effects on our main conclusions (see *Materials and Methods: Eye movements*).

### Foveal V4 is more sensitive to fine-scale scrambling than peripheral V4

We next looked at how scrambling sensitivity changed with visual eccentricity. We split our neural population into 4 nonoverlapping sets that each covered distinct ranges of eccentricity. Median group eccentricities ranged from 1.5 degrees to 9.2 degrees. We then measured within-family discriminabilities within these eccentricity subpopulations (Figure 5A).

**Figure 5:**
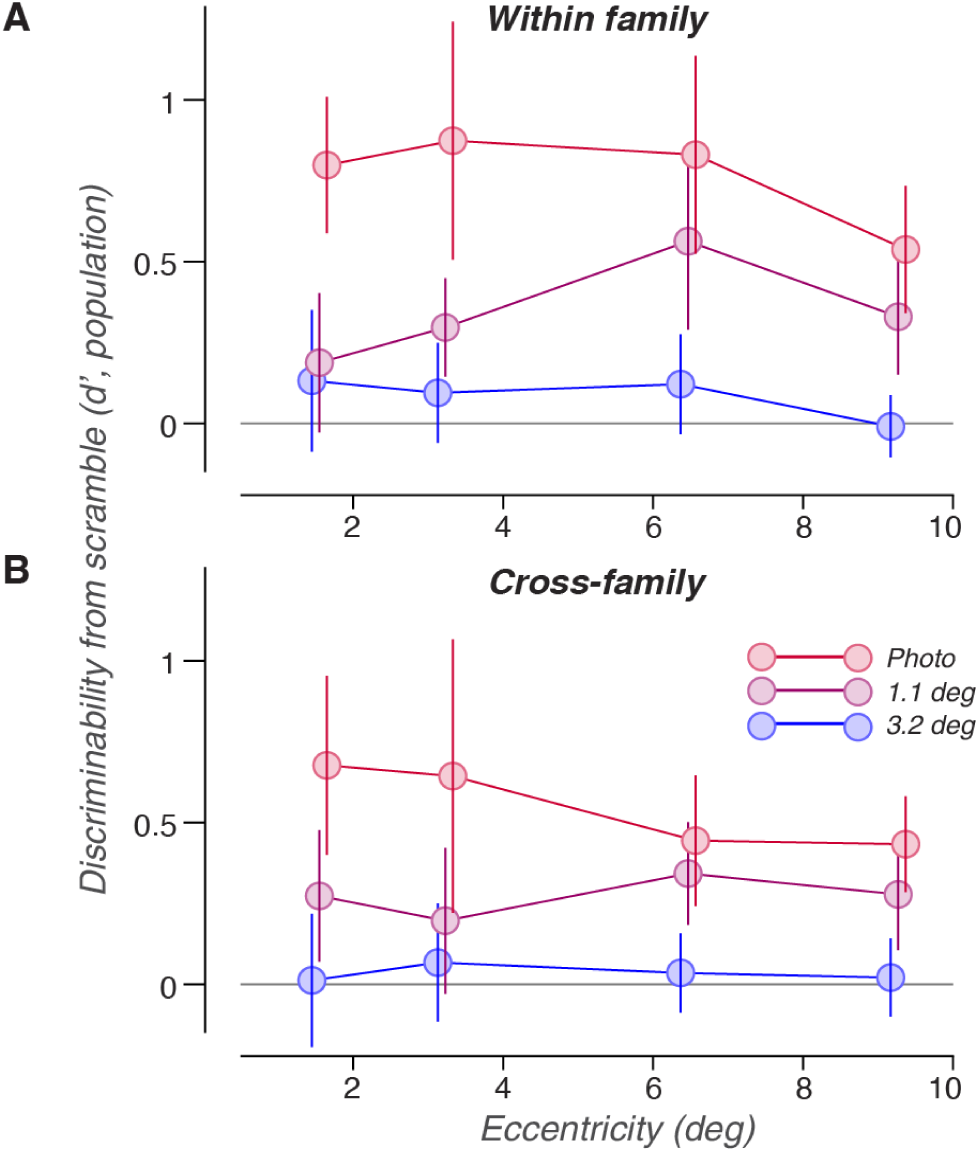
Foveal V4 is more sensitive to fine-scale scrambling than peripheral V4. A) Within-family discriminability between fully scrambled image responses and either photographic images (red) or partially scrambled images (purple, blue), using responses from four distinct groups of subpopulations whose receptive field centers cover different ranges of eccentricity. Discriminabilities were computed for subpopulations of 18 neurons. Discriminations for different pooling region sizes are denoted by different colors. B) Cross-family discriminability (as in Figure 4B), for four groups of subpopulations with non-overlapping eccentricities.

Models of spatial pooling predict that neurons spatially average information within their receptive fields, and that the spatial scale of this process increases with increasing eccentricity (Freeman and Simoncelli, 2011; Rosenholtz et al., 2019). These models predict that foveal V4 neurons average within small regions and thus preserve information about fine details. As such, they should be strongly disrupted by all scrambling conditions, even those with small pooling regions (1.1 deg diameter). Conversely, peripheral V4 responses average over larger regions and thus lose information about fine details. That is, peripheral V4 responses are already, in some sense, “scrambled,” and should thus be relatively invariant to scrambling within small pooling regions.

Consider the differences between the curves in Figure 5A. For foveal populations, the spatial pooling framework predicts that the difference in discriminability between photographic and 1.1 degree pooling regions should be *large* (Figure 5A, left). Conversely, for peripheral populations, the difference in discriminability between photographic and 1.1 degrees should be *small* (Figure 5A, right). This is the trend that we observed. While the difference between photographic and 1.1 degrees at all eccentricities were substantial, they were typically larger for foveal populations (Δd’=0.550 between 1.1 degree and photographic) and smaller for peripheral populations (Δd’=0.252). Across all subpopulations, this difference in discriminability decreased 0.047 Δd’/degree moving from foveal to peripheral receptive fields.

We extended the eccentricity analysis to cross-family decoders (as in Figure 4A, grey points), and again found a difference between foveal and peripheral responses (Figure 5B). Foveal population responses were more disrupted by scrambling with small pooling regions (Δd’=0.4, 1.1 degree vs. photographic) and peripheral population responses were less disrupted (Δd’=0.16, overall Δd’ decreased 0.043/degree from fovea to periphery).

We draw two primary conclusions. First, foveal responses were more strongly degraded by partial scrambling than peripheral responses, consistent with predictions from spatial pooling models (Freeman and Simoncelli, 2011; Rosenholtz et al., 2019; Ziemba and Simoncelli, 2021) and human perception (Pelli and Tillman, 2008). Second, V4 populations at all eccentricities were substantially affected by scrambling, even when that scrambling was perceptually subtle and local (confined to subregions of ∼1 degree diameter), and even for subpopulations with peripheral receptive fields (∼9 degrees eccentricity).

### V4 responses discriminate photographic images better than scrambled images

Having established that V4 populations could detect the presence or absence of photographic structure, we next turned to a different comparison: whether V4 responses could discriminate between pairs of image families, and whether that discrimination was affected by image scrambling. Distinguishing between photographic images is a task that has frequently been used to characterize late ventral stream processing (Rust and DiCarlo, 2010; Yamins et al., 2014; Majaj et al., 2015; Fyall et al., 2017). Comparing photographic image discrimination to the analogous task of scrambled image discrimination allows us to quantify the extent to which discriminability depends on image statistics (features shared between scrambled and photographic images) or on the spatial configuration of image features (which are only present in photographic images).

We first measured family discriminability for individual V4 neurons by averaging d’ values over all possible pairs of families. Single-neuron discriminability (Figure 6A) was systematically higher for responses to photographic images than for responses to scrambled images (median photographic d’=0.43, vs median scramble d’=0.37).

**Figure 6:**
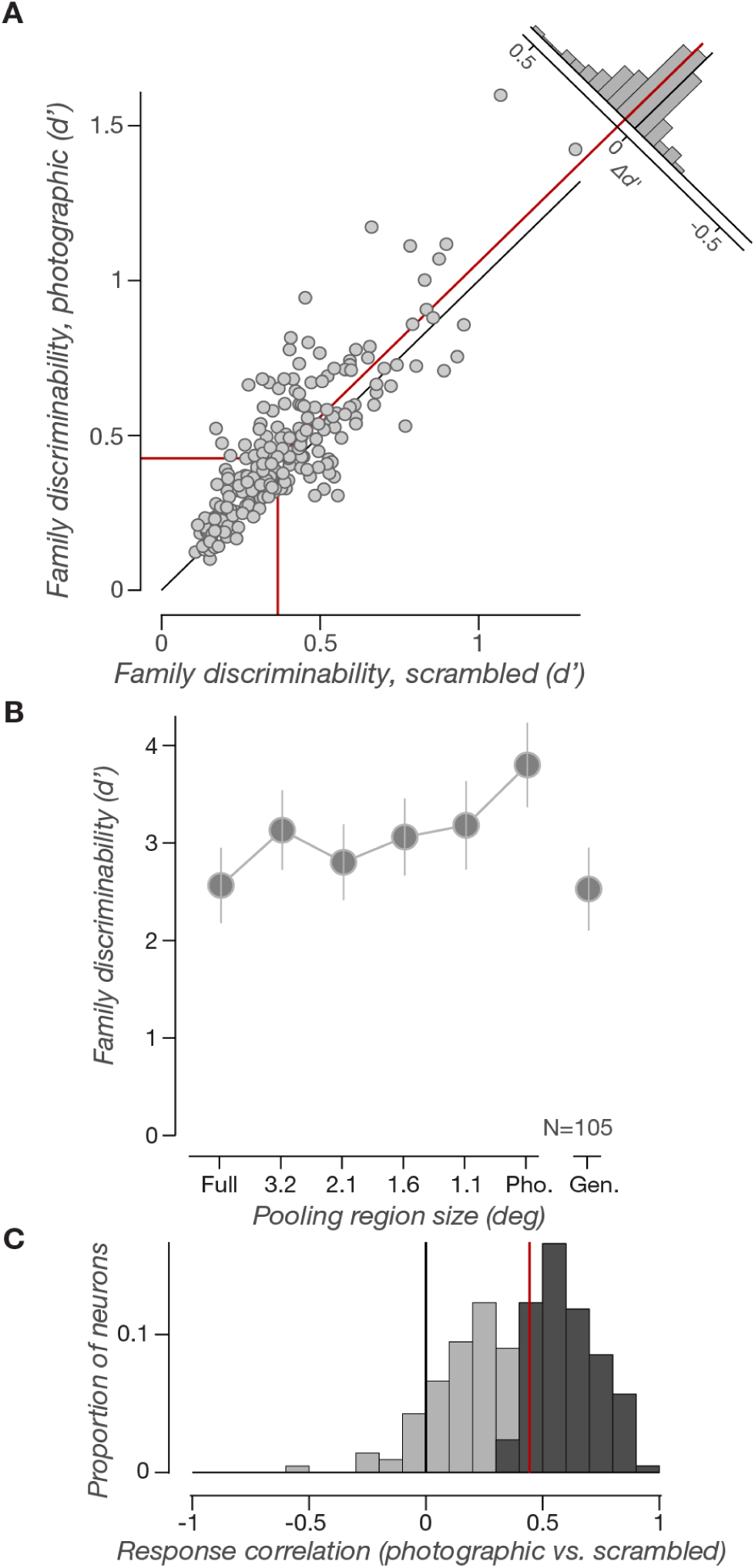
V4 discriminates photographic scenes more reliably than their scrambled counterparts. A) Single neuron family discriminability for photographic images, plotted against family discriminability for scrambled texture images. Discriminabilities were measured as the median d’ across all possible pairs of image families. The inset above shows the distribution over neurons of the difference between photographic and scrambled d’. Red lines denote mean values for the photographic distribution (d’=0.43), the scrambled distribution (d’=0.37), and their difference (Δd’=0.06). A majority of neurons (151/211) show higher discriminability between photographic images than scrambled images. B) Average neuronal population family discriminability for full and intermediate levels of scrambling, and for photographs (N=105 neurons). Error bars show standard deviations across resampled neuronal populations. The far right point (“Gen.”) shows the discriminability of a generalization classifier trained on scrambled images and tested on photographic images. Both the full and generalization conditions are ∼2/3 the discriminability of the photograph condition. C) Distribution of individual neuron correlations between firing rate responses to photographic images and to matched scrambled images. Neurons with correlations significantly greater than 0 (p<0.05) are in dark grey (122/211). The red line denotes the median (r=0.443).

Next, we measured discriminability for V4 population responses. For each pair of image families, we used LDA to find neuronal weights that best separated the two sets of responses (*Materials and Methods: Classification with LDA*). This LDA was cross-validated across distinct shifts within each image family (as in Figure 1A). As such, this analysis measures “translation-invariant” discriminability, akin to previous measurements of “position invariant” object recognition performance in V4 (Rust and DiCarlo, 2010; El-Shamayleh and Pasupathy, 2016).

Previous work has shown that populations of V4 neurons can better classify images of photographs than images of matched textures (Rust and DiCarlo, 2010). Consistent with those results, we found that population discriminability for scrambled image responses was ∼2/3 as large as for photographic responses (Figure 6B). We also observed a trend similar to the photographic vs. scrambled image task (Figure 4A), where adding a small amount of scrambling to photographic images led to a large change in discriminability. The gap between photographic images and their closest, least scrambled counterparts (1.1 deg. scrambling regions) was comparable in size to the gap between those least scrambled images and the fully scrambled images (Δd’=0.619 vs. Δd’=0.616, respectively).

To investigate whether family discrimination performance was affected by receptive field eccentricity (as in Figure 4), we again split our population into 4 non-overlapping subpopulations covering distinct eccentricities. As before, we found that partial scrambling more strongly degraded discrimination in foveal populations than in peripheral populations (foveal Δd’= 0.36, 1.1 deg vs. photographic, peripheral=0.28, decreased 0.015 Δd’/degree from fovea to the periphery).

Finally, we investigated how the structure of the V4 population response changed for scrambled and photographic images. Scrambled and photographic images contain the same textural features, but only photographic images have “object-like” structure. To determine how adding object-like features changed population responses, we trained a generalization classifier: a classifier trained on scrambled image responses and tested on photographic image responses.

The relative performance of this generalization classifier will depend on the geometry of the V4 population response to scrambled and photographic images. One possibility is that photographic images drive the same pattern of activity as scrambled images, but increase the response gain, leading to stronger responses and better discriminability. If this is the case, generalization discriminability should be as large as photographic image discriminability. A second possibility is that the textural features of scrambled images are encoded using patterns of activity that are fully independent of (orthogonal to) the patterns driven by photographic image features. In this case, generalization discriminability should be similar to scrambled image discriminability. A final possibility is that photographic features could supersede and actively suppress the representation of textural features. In this case, photographic features would not only be encoded by independent patterns of activity, but would also reduce or eliminate response gain within the patterns of activity representing textural features. Thus, generalization discrimination would be lower in magnitude than scrambled image discriminability.

The discriminability of this generalization classifier (Figure 6B, “Gen.”) was similar to that of the scrambled image classifier, and roughly 2/3 that of the photographic classifier. This is most consistent with the second possibility discussed above: that V4’s representation of “object-like” features is encoded in neuronal dimensions orthogonal to those representing textural features. These results also suggest that the representation of textural features is not suppressed by the presence of object-like features. Rather, textural features are encoded independently of object-like features, and are similarly informative for both scrambled and photographic image responses. Lastly, these results suggest that most of the family discrimination signal (2/3) is attributable to textural feature responses, and that only 1/3 is attributable to object-like features.

### A model of perceptual similarity predicts V4 responses to image scrambling

For both comparisons discussed thus far (photographic structure detection and image family discrimination) we observed that photographic image discriminability was strongly reduced by small amounts of image scrambling. This surprised us, as we chose these 6 levels of scrambling to correspond to roughly equal perceptual increments (Figure 1C). By inspection, examples from the least scrambled condition (pooling size 1.1 deg, see Figure 1C) appear quite similar to the original photographs. Nonetheless, classification performance differed strongly between the two cases. We sought to compare these disparate neural and perceptual effects.

We created a neural distance metric by building normalized population response vectors for each image, then measuring the Euclidean distance between pairs of image vectors (see *Materials and Methods: Perceptual and neural distance metrics*). On average, neural distances were larger between photographic images and any of their scrambled counterparts than they were between any two scrambled conditions (Figure 7A). In the plot in Figure 7A, this is visually evident as lighter cell values on the left column and bottom row.

**Figure 7:**
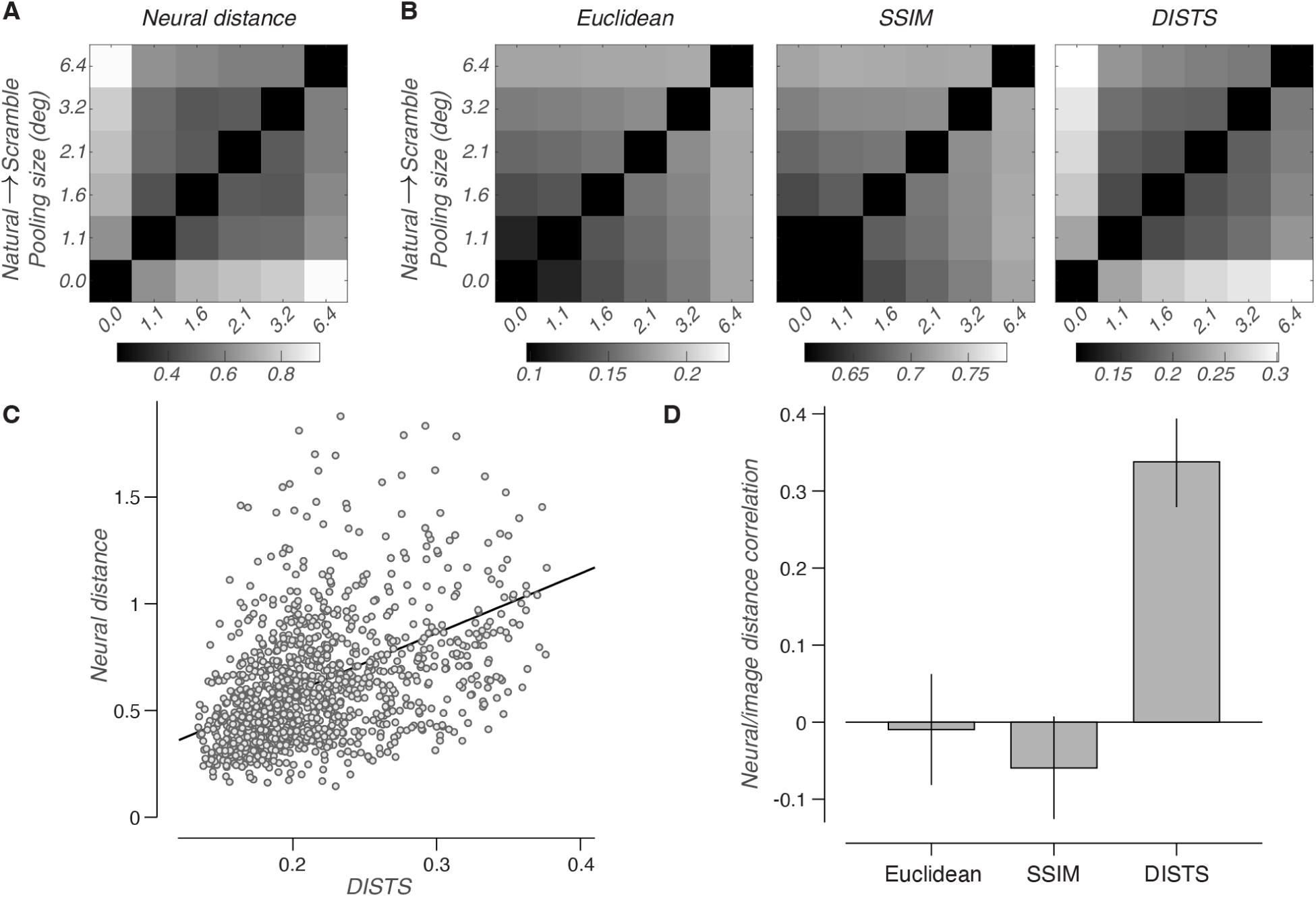
A model of perceptual discriminability (DISTS) predicts V4 population responses. A) Neural distances, measured between pairs of V4 population responses. Distances were only measured between matched images (same family and shift) with different levels of scrambling. Pooling regions range from fully scrambled images (6.4 degrees) to photographic images (0.0 degrees). Shading indicates Euclidean distance between pairs of normalized population vectors, averaged over all appropriate image pairs. The white banding on the left and bottom edges indicates that neural populations categorically separate photographic images from all scrambled conditions. B) Image similarity, measured between matched images with different levels of scrambling. We used three different distance metrics: 1) average RMS pixel distance (left), 2) SSIM (center), and 3) DISTS (right). Pixel distance and SSIM differ from neural distance, while DISTS appears similar. C) Neural distance vs. DISTS for each measured pair of images. For 20 image families and 4 shift conditions per family, we measured 30 image comparisons (2400 total pairs plotted). The linear best fit is plotted in black. D) Correlations between neural distances and the three image similarity metrics, averaged over resampled neural subpopulations (N=105 neurons). Error bars represent standard deviations across subpopulations. DISTS predicts neural distance, while pixel distance and SSIM do not.

Next, we asked whether pixel-based measurements of the images would show a similar separation of photographs from scrambled images. We considered three distance metrics: 1) pixel-based Euclidean distance, 2) the Structural Similarity index (SSIM (Wang et al., 2004)), and 3) Deep Image Structure and Texture Similarity (DISTS (Ding et al., 2022)). Both SSIM and DISTS are designed to quantify human perceptual judgments of image distortion. However, DISTS design has two important differences from pixel distance and SSIM. First, DISTS is robust to image distortions that preserve texture identity. Second, the free parameters in DISTS were directly fit to predict human judgements of image similarity for distorted and photographic images. Because of these two differences, DISTS better predicts perceptual similarity ratings than either Euclidean distance or SSIM.

For both pixel distance and SSIM, we observed that image distances (averaged over image pairs) displayed a smooth gradation from fully scrambled images to partially scrambled images to photographic images (Figure 7B, left and center). This did not match the pattern observed in neural distance. Conversely, the DISTS metric revealed a sharp distinction between photographic images and all scrambled conditions (Figure 7B, right) that more closely resembled neural distance. Across all individual pairs of images, we found that distances measured with DISTS were significantly correlated with V4 neural distances (Figure 7C-D), while measurements using pixel distances or SSIM showed essentially no correlation (Figure 7D). This suggests that the large difference between V4 responses to photographs and all scrambled conditions may reflect the sensitivity of human observers to small, unnatural distortions in photographic images (Wallis et al., 2019; Broderick et al., 2025).

### Sensitivity to image scrambling emerges slowly

To characterize how V4’s sensitivity to image scrambling changed over time, we trained classifiers for both the photographic structure detection task and the image family discrimination task in a series of 50 ms bins that spanned the temporal extent of the response.

First, we examined the time course of photographic structure detection. Specifically, we measured discriminability for the combined set of all image families (“cross-family,” as in Figure 4A), comparing fully scrambled images to either photographic images or partially scrambled images (Figure 8A). Sensitivity was slightly lower in the smaller time bins compared with the full temporal extent of the response (d’=1.33 peak 50 ms bin, vs. d’=1.48 for the full 350 ms response), but still exhibited a large difference between photographic image sensitivity (red curve) and sensitivity for intermediate levels of scrambling (maroon/blue curves). For the three conditions for which we could robustly extract a time course, latencies to half-maximum performance were similar (photographic images: 99±11.4 ms, 1.07 deg.=103±10.6 ms, 1.60 deg.=122±4.7 ms).

**Figure 8:**
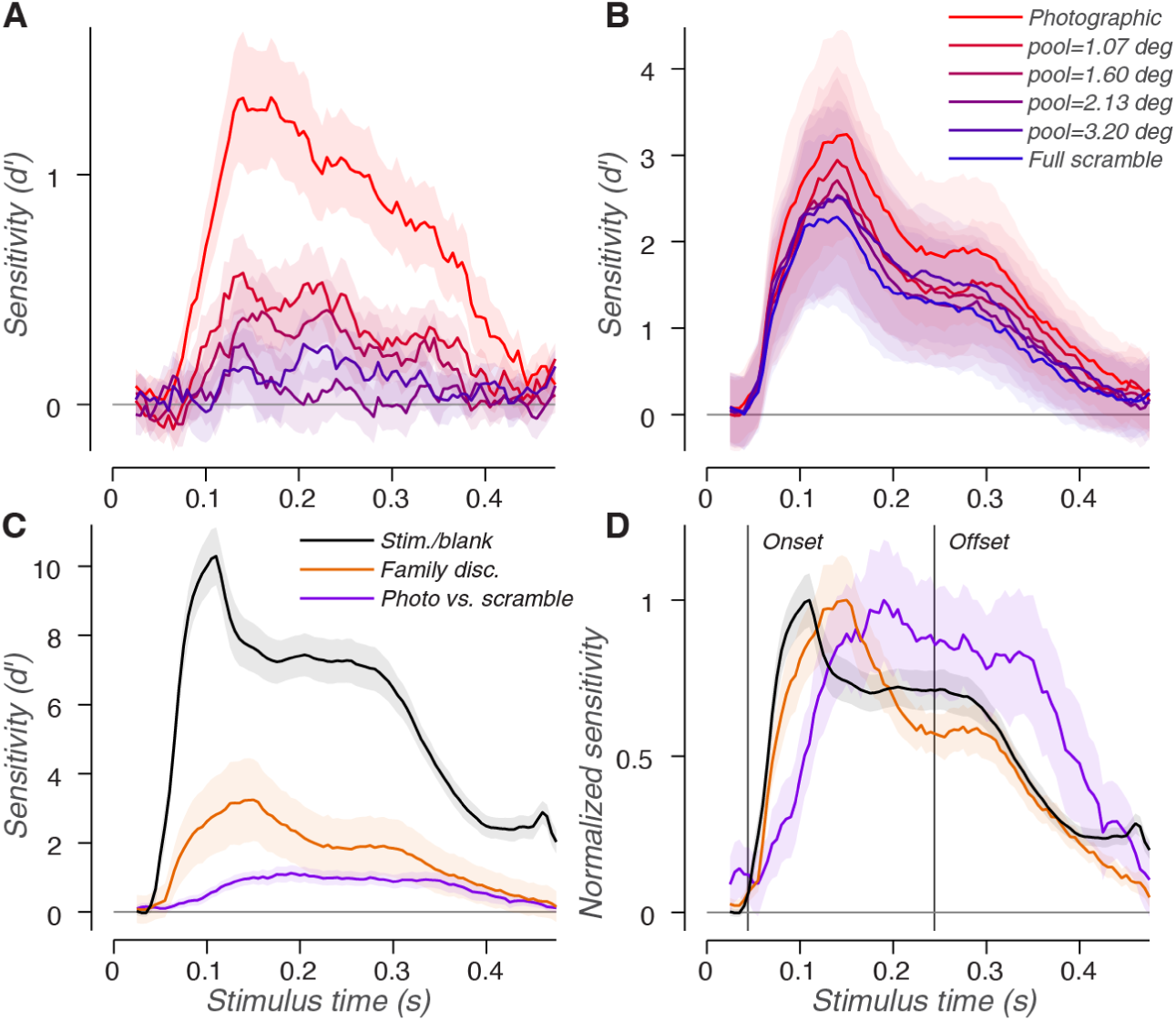
Sensitivities for different tasks emerge at different times. A) Discriminability between fully scrambled images and either photographic images (red) or intermediately scrambled images (other colors), plotted over time. Sensitivities were computed within 50 ms windows. The shaded regions represent standard deviations across subpopulations. B) Pairwise image family discriminability plotted over time. As in A), responses were calculated in 50 ms bins. Plots of different colors represent discriminations between photographic images (red), between fully scrambled images (blue), or between intermediately scrambled images (other colors). Shaded regions represent standard deviations across subpopulations. C) Comparison of sensitivity time courses for 3 different tasks: discrimination between image trials and blank trials (black), pairwise photographic family discrimination (orange), and photographic structure detection (purple). D) The 3 task traces from C) replotted as normalized sensitivities, such that peak sensitivity is 1. Different task sensitivities emerge at different times. Vertical lines mark stimulus/blank discriminability latencies for onset (44 ms) and offset (244 ms).

Next, we looked at the dynamics of pairwise image family discriminability (Figure 8B). The peak sensitivity across all 50 ms bins was only slightly lower than the sensitivity calculated using the full 350 ms extent of the response (d’=3.24 peak 50 ms bin, vs. d’=3.80 full 350 ms). At early times (∼150 ms, near peak performance), discriminabilities showed a graded improvement between fully scrambled images, intermediate levels of scrambling, and photographic images. At later time points (∼300 ms), discriminabilities for full and intermediate levels of scrambling became similar, while photographic image discriminability remained well separated from the intermediate scrambling conditions. Despite peaking at different sensitivities, in all 6 conditions, discriminability took similar amounts of time to reach half-maximum (66–77 ms).

We next compared the time courses of these tasks. In addition to photographic structure detection (Figure 8A) and image family discrimination (Figure 8B), we also measured the time course of discriminability between stimulus and blank trials. This can be thought of as a baseline measurement of V4’s response to large changes in contrast. Discriminabilities for these three tasks strongly differed in magnitude (Figure 8C). Stimulus-blank discriminability was more than twice as large as image family discriminability, which was again much larger than photograph vs. scramble discriminability. To compare timings we computed *normalized sensitivity*: each time course divided by its peak sensitivity (Figure 8D). The normalized curves show that stimulus-blank discriminability emerges first (half-max latency=66±2.1 ms), followed closely by the photographic family discrimination task (71±4.9 ms). Photographic vs. scrambled sensitivity emerges last, with substantial delay (99±8.6 ms).

We estimated that stimulus/blank discriminability first began rising in an analysis bin centered at a latency of 44 ms. Because the classification analysis is performed in 50 ms wide bins, this implies that V4 responses begin to rise at the edge of this bin, 25 ms later, at 69 ms. This measure of V4’s onset latency is consistent with previous reports (Lee et al., 2007; Oleskiw et al., 2018; Zamarashkina et al., 2020; Kim et al., 2022; Rodríguez-Deliz et al., 2025). We defined the “offset latency” as a time bin centered 200 ms after this onset latency, at 244 ms. Activity after this point represents V4’s response to the removal of the image from the screen. In all cases, we observed both robust firing rates and positive classification sensitivities for at least 150 ms after this offset point. Notably, of the three tasks, the photographic vs. scramble task sensitivities persisted at near-peak values up to and beyond the stimulus offset latency. The fact that this task was characterized by both a slow onset latency and slow decay suggests that the “cross-family” photograph vs. scramble signal may arise from recurrent or feedback mechanisms.

## Discussion

We created families of images that are matched in their local texture, and used them to show that population responses in V4 can reliably distinguish images of objects from images of textures. This object selective signal was strongest when spatial feature scrambling was constrained within roughly 1.5 degrees. Neural discriminability resembles human perceptual discriminability (as simulated by an image similarity metric), emerges slowly, and persists through the temporal extent of the response.

### Distinguishing objects from textures

As in previous reports, V4 neuronal selectivities spanned a continuum between those preferring objects and those preferring texture (Kim et al., 2019; Willeke et al., 2023). We found a relatively balanced distribution of these two populations, so that photographic images and scrambled images drove similarly strong responses when averaged across the whole V4 population. While response magnitude did not differentiate photographic and scrambled images, response variance did. V4 neurons responded with a wider dynamic range to photographic images than to scrambled images, which may explain why photographic image responses are more discriminable than scrambled responses (Rust and DiCarlo, 2010).

V4 population responses distinguish objects from textures. These responses are then fed forward to area IT, where the representation of the distinction between objects and textures grows stronger (Rust and DiCarlo, 2010), and where texture responses are suppressed (Lee et al., 2024). Our results draw similarities to psychophysical studies of crowding, which have shown that crowded visual scenes, like textures, are encoded with a compressed perceptual representation (Bouma, 1970; Levi, 2008; Pelli and Tillman, 2008; Balas et al., 2009; Ziemba and Simoncelli, 2021). V4 is a critical stage of this perceptual process: human observers with more cortical area devoted to V4 are less likely to perceive objects as crowded within their surrounding texture (Kurzawski et al., 2024). V4 may be where the distinction between objects and textures first emerges in an object-invariant manner.

Some previous studies comparing object and texture responses have shown results that seem to differ from ours. First, two human fMRI studies have reported a stronger V4 response to photographic images than scrambled images (Movshon and Simoncelli, 2014; Long et al., 2018). This may be due to differences between single unit neurophysiology and fMRI. While the hemodynamic responses captured by BOLD fMRI are related to pooled neural activity, this relationship is not linear (Logothetis et al., 2001). In our data, V4 neural responses had a greater dynamic range for photographic images than for scrambled images, without a change in the mean. A nonlinear relationship between neuronal firing and BOLD could convert this difference in range into a difference in the mean fMRI signal.

Second, two studies using an image synthesis method similar to ours have reported that original images drive stronger neural responses than scrambled images (Long et al., 2018; Kramer et al., 2023). One possible reason for this difference with our study is that we used photographs with natural backgrounds, while these studies used “TexForms” that were cropped and placed on grey backgrounds. The backgrounds surrounding objects in our images may have suppressed V4 responses, relative to those measured with TexForms.

Finally, a study using a texture synthesis model based on a deep neural network (DNN) found that fMRI responses across the ventral stream, including areas V4 and IT, could not reliably distinguish between their photographic images and scrambled counterparts (Jagadeesh and Gardner, 2022). While these results appear inconsistent with ours, they are based on texture stimuli whose generation was constrained by statistics from the deepest layers of a DNN. These deep layers can predict firing rate responses in later visual areas like IT (Yamins et al., 2014). Conversely, the statistics of the Portilla-Simoncelli texture model are primarily correlations of V1-like statistics, which are more naturally associated with area V2 (Portilla and Simoncelli, 2000; Freeman et al., 2013), or V4 (Okazawa et al., 2016). The DNN-based statistics capture more characteristically “object-like” features for synthesis, and therefore create images sets that are less strongly differentiated from photographs than the image sets used in this study.

### Object and texture discrimination

Object recognition is one of the primary computational goals of the ventral stream of visual cortex (DiCarlo et al., 2012). V4 responses can distinguish between images of objects, and only 1/3 of this object recognition signal relied on object-specific features in the images, while 2/3 relied on properties that were shared with the textures (Figure 4A). That is, simple, V1-like image features (like spatial frequency and orientation) and more complex, V2-like “naturalistic” features (Freeman et al., 2013; Okazawa et al., 2015, 2016; Lee et al., 2024) carry most of the object recognition signal in V4. This ratio likely reflects that V4 is an early stage of object selectivity. Comparable measurements in downstream area IT have demonstrated an object recognition signal that is more strongly dependent on object-specific features (Rust and DiCarlo, 2010).

### Responses to partially scrambled images

Neural responses to partially scrambled images were more similar to fully scrambled responses than to photographic responses, even when scrambling was perceptually subtle. This is consistent with previous observations that V4 responses are robustly driven and modulated by images of shapes and objects (Pasupathy et al., 2020). Neuronal responses in V4 are selective for complex shape (Kobatake and Tanaka, 1994; Pasupathy and Connor, 1999, 2001) and lesions to area V4 profoundly disrupt form-processing performance (Merigan, 1996). Many neurons in V4 are selective for the sharpness of object edges (Oleskiw et al., 2018) and monocular cues for three-dimensional shape (Srinath et al., 2021), both of which are disrupted by image scrambling. Similarly, human object perception is highly sensitive to disruptions in boundary closure (Elder et al., 2018) and to scrambling (Wallis et al., 2019; Broderick et al., 2025). Inspired by computer vision research, we found that DISTS (a model of perceived image distortion (Lin et al., 2019; Ding et al., 2022)) predicted this perceptual sensitivity to image scrambling and could account for V4’s sensitivity to scrambling.

### Response dynamics

V4’s response to natural images evolves with time. We found that V4 responses quickly discriminate between different object images (∼10 ms after stimulus onset), suggesting that the object selectivity signal is primarily built from feedforward signals originating in areas upstream. This is consistent with other stimulus-driven properties that emerge quickly, such as selectivity for complex contours (Yau et al., 2013) or shape (Oleskiw et al., 2018). Conversely, selectivity between photographic and scrambled images emerges later (∼50 ms after stimulus onset), and grows at a slower rate than object selectivity. This is similar to the time course for V4’s selectivity to perceptually-salient textural features (Kim et al., 2022). And all of these stimulus-driven properties emerge faster than more “cognitive” processes, such as attentional modulation (Motter, 1994) and the “filling in” of occluded stimuli (Fyall et al., 2017).

The relatively delayed time course of modulation by the degree of image scrambling suggests that selectivity for object-specific image features might rely on recurrent processing within V4, or on feedback from downstream ventral areas, such as posterior inferotemporal cortex (PIT) (Felleman and Van Essen, 1991), where object images modulate responses more strongly than matched texture images (Rust and DiCarlo, 2010). Such feedback signals may propagate further upstream than V4 – it has recently been reported that some neurons in V1 show a preference for photographic images over scrambled images (Chen et al., 2022; Kramer et al., 2023).

### Spatial pooling

Many perceptual phenomena provide evidence that spatial pooling of image statistics is a characteristic of ventral stream processing (Lettvin, 1976; Wilkinson et al., 1997; Parkes et al., 2001; Pelli et al., 2004; Pelli and Tillman, 2008; Balas et al., 2009; Greenwood et al., 2009; Freeman and Simoncelli, 2011; Rosenholtz et al., 2019; Ziemba and Simoncelli, 2021). Models of pooling often associate the spatial extent of pooling with receptive field sizes in ventral stream areas. Here, we estimated V4’s critical distance for spatial pooling to be roughly 1.5 degrees (Figure 4A). This is much smaller than the typical receptive field sizes of V4 neurons (Gattass et al., 1988), and is more consistent with receptive field sizes in upstream areas like V2 (Gattass et al., 1981). This is also supported by psychophysical evidence: image scrambling is most difficult to perceive when it is confined within the spatial extent of V2 receptive fields (Freeman and Simoncelli, 2011; Ziemba and Simoncelli, 2021). These results suggest that V2 may be the last processing stage in the ventral stream to represent *only* local image statistics – stuff – and that V4, and presumably areas downstream, compute a representation that more explicitly captures visual objects – things.

## Acknowledgements

We are grateful to Kahlia Gronthos, Sullivan Bacerdo, and Kaitlyn Holman for their assistance, and to Manu Raghavan for his help with hardware and software. This work was supported by grants from the National Institutes of Health (EY022428) and the Simons Foundation (543019) to J.A.M. and E.P.S, as well as the Flatiron Institute of the Simons Foundation (T.D.O. and E.P.S.). J.D.L was supported in part by a Leon Levy Fellowship in Neuroscience. Computational support was provided through NYU IT High Performance Computing services.

## Author contributions

Conceptualization and methodology, J.D.L, E.P.S., and J.A.M.; Investigation, J.D.L., T.D.O., and L.P.; Writing, J.D.L., E.P.S., and J.A.M.

